# Heterozygous variants in KMT2E cause a spectrum of neurodevelopmental disorders and epilepsy

**DOI:** 10.1101/566091

**Authors:** Anne H O’Donnell-Luria, Lynn S Pais, Víctor Faundes, Jordan C Wood, Abigail Sveden, Victor Luria, Rami Abou Jamra, Andrea Accogli, Kimberly Amburgey, Britt Marie Anderlid, Silvia Azzarello-Burri, Alice A Basinger, Claudia Bianchini, Lynne M Bird, Rebecca Buchert, Wilfrid Carre, Sophia Ceulemans, Perrine Charles, Helen Cox, Lisa Culliton, Aurora Currò, Deciphering Developmental Disorders (DDD) Study, Florence Demurger, James J Dowling, Benedicte Duban-Bedu, Christele Dubourg, Luis F Escobar, Alessandra Ferrarini, Tobias B Haack, Mona Hashim, Solveig Heide, Katherine L Helbig, Ingo Helbig, Raul Heredia, Delphine Heron, Bertrand Isidor, Amy R Jonasson, Pascal Joset, Boris Keren, Fernando Kok, Hester Y Kroes, Alinoë Lavillaureix, Xin Lu, Saskia M Maas, Gustavo HB Maegawa, Carlo LM Marcelis, Saga Elise Eiset, Paul R Mark, Mercelo R Masruha, Heather M McLaughlin, Kirsty McWalter, Esther U Melchinger, Saadet Mercimek-Andrews, Caroline Nava, Manuela Pendziwiat, Richard Person, Gian Paolo Ramelli, Luiza LP Ramos, Anita Rauch, Caitlin Reavey, Alessandra Renieri, Angelika Rieß, Amarilis Sanchez-Valle, Shifteh Sattar, Carol Saunders, Niklas Schwarz, Thomas Smol, Myriam Srour, Katharina Steindl, Steffen Syrbe, Jenny C Taylor, Aida Telegrafi, Isabelle Thiffault, Doris A Trauner, Helio van der Linden, Silvana van Koningsbruggen, Laurent Villard, Ida Vogel, Julie Vogt, Yvonne G Weber, Ingrid M Wentzensen, Elysa Widjaja, Jaroslav Zak, Samantha Baxter, Siddharth Banka, Lance H Rodan

**Affiliations:** Division of Genetics and Genomics, Boston Children’s Hospital, Harvard Medical School, Boston, MA 02115, USA.; Broad Center for Mendelian Genomics, Broad Institute of MIT and Harvard, Cambridge, MA 02142, USA.; Division of Evolution & Genomic Sciences, School of Biological Sciences, Faculty of Biology, Medicine and Health, University of Manchester, Manchester M13 9PL, UK.; Laboratorio de Genética y Enfermedades Metabólicas, Instituto de Nutrición y Tecnología de los Alimentos (INTA), Universidad de Chile, Santiago, Chile.; Department of Systems Biology, Harvard University Medical School, Boston, MA 02115, USA.; Institute of Human Genetics, University of Leipzig Medical Center, Leipzig 04103, Germany.; Department of Pediatrics, Department of Neurology and Neurosurgery, McGill University, Montreal QC H4A 3J1, Canada.; DINOGMI - Università degli studi di Genova, 16126 Genova, Italy.; UOC Neurochirurgia, IRCCS Istituto Gaslini, 16147 Genova, Italy.; Division of Neurology, Department of Pediatrics, Hospital for Sick Children, University of Toronto, Toronto M5G 2C4, Ontario, Canada.; Department of Molecular Medicine and Surgery, Centre for Molecular Medicine, Karolinska Institutet and Department of Clinical Genetic, Karolinska University Hospital, Stockholm 141 86, Sweden.; Institute of Medical Genetics, University of Zurich, Schlieren-Zurich CH-8952, Switzerland.; Neuroscience Center Zurich, University of Zurich and ETH, Zurich 8057, Switzerland.; Genetics, Cook Children’s Physician Network, Fort Wort, TX 76104, USA.; Pediatric Neurology, Neurogenetics and Neurobiology, Unit and Laboratories, Neuroscience Department, A Meyer Children’s Hospital, University of Florence, 50139 Florence, Italy.; Department of Pediatrics, University of California, San Diego, San Diego, CA 92093, USA.; Division of Genetics, Rady Children’s Hospital of San Diego, San Diego, CA 92123, USA.; Institute of Medical Genetics and Applied Genomics, University of Tuebingen, Tuebingen 72074, Germany.; Laboratoire de Génétique Moléculaire et Génomique, CHU de Rennes, Rennes 35033, France.; Department of Genetics, Pitié-Salpêtrière Hospital, Assistance Publique - Hôpitaux de Paris, Paris 75013, France.; GRC Déficience Intellectuelle et Autisme, Sorbonne University, Paris 75006, France.; West Midlands Regional Clinical Genetics Service and Birmingham Health Partners, Birmingham Women’s and Children’s Hospital NHS Foundation Trust, Birmingham B15 2TG, UK.; Department of Neurology, Children’s Mercy Hospital and Clinics, MO 64108, USA.; Medical Genetics, University of Siena, 53100 Siena, Italy.; Genetica Medica, Azienda Ospedaliera Universitaria Senese, 53100 Siena, Italy.; Wellcome Sanger Institute, Wellcome Genome Campus, Hinxton CB10 1SA, UK.; Service de Génétique Clinique, Centre de Référence Maladies Rares CLAD-Ouest, CHU de Rennes, 35033 Rennes, France.; Centre de Génétique Chromosomique, GHICL Hôpital Saint Vincent de Paul, 59020 Lille, France.; Faculté de médecine de l’UCL, Université Catholoique de Lille, 59800 Lille, France.; Laboratoire de Génétique Moléculaire & Génomique, CHU de Rennes, 35033 Rennes, France.; St. Vincent’s Childrens Hospital, Indianapolis, IN 46260, USA.; Medical Genetic Unit, Italian Hospital of Lugano, Lugano, Switzerland; Università della Svizzera Italiana, 6900 Lugano, Switzerland.; Oxford NIHR Biomedical Research Centre, Wellcome Centre for Human Genetics, University of Oxford, Oxford OX3 7BN, UK.; Division of Neurology and Department of Biomedical and Health Informatics (DBHi), Children’s Hospital of Philadelphia, PA 19104, USA.; Department of Neurology, University of Pennsylvania, Perelman School of Medicine, Philadelphia, PA, 19104 USA.; Department of Neuropediatrics, Christian-Albrechts-University of Kiel, 24105 Kiel, Germany.; GeneDx, Gaithersburg, MD 20877, USA.; Department of Genetics, Centre de Référence Déficiences Intellectuelles de Causes Rares, 75012 Paris, France.; GRC Déficience Intellectuelle et Autisme, Sorbonne University, 7500 Paris, France.; Service de Génétique Médicale, Hôpital Hôtel-Dieu, CHU de Nantes, 44093 Nantes France.; Division of Genetics and Metabolism, Department of Pediatrics, University of Florida, FL 32610, USA.; Mendelics Genomic Analysis, Sao Paulo, Brazil 04013.; Department of Medical Genetics, University Medical Center Utrecht, 3584 CX Utrecht, Netherlands.; Ludwig Institute for Cancer Research, Nuffield Department of Clinical Medicine, University of Oxford, Oxford OX3 7DQ, UK.; Amsterdam UMC, University of Amsterdam, Department of Clinical Genetics, 1012 WX Amsterdam, The Netherlands.; Department of Clinical Genetics, Radboud University Medical Centre, 6525 GA Nijmegen, Netherlands.; Department of Clinical Genetics, Aarhus University Hospital, 8200 Aarhus, Denmark.; Spectrum Health Medical Genetics, Grand Rapids, MI 49544, USA.; Department of Neurology and Neurosurgery, Universidade de Federal de São Paulo, São Paulo 04023, Brazil.; Division of Clinical and Metabolic Genetics, Department of Pediatrics, University of Toronto, The Hospital for Sick Children, Toronto, Ontario, M5G 1X8, Canada.; Department of Neuropediatrics, University Medical Center, Schleswig Holstein, 24105 Kiel, Germany.; Pediatric Department of Southern Switzerland, Neuropediatric Unit, San Giovanni Hospital, 6500 Bellinzona, Switzerland.; Rare Disease Initiative Zürich, Clinical Research Priority Program for Rare Diseases, University of Zurich, CH-8006 Zurich, Switzerland.; Department of Pediatrics, Division of Genetics and Metabolism, University of South Florida, Tampa, FL 32610, USA.; Section of Pediatric Neurology, Rady Children’s Hospital, San Diego, CA 92123, USA.; Departments of Neurosciences and Pediatrics, University of California San Diego, La Jolla, CA 92093, USA.; Center for Pediatric Genomic Medicine, Children’s Mercy Hospital and Clinics, Kansas City, MO 64108, USA.; School of Medicine, University of Missouri, Kansas City, Mo 64108, USA.; Department of Neurology and Epileptology, Hertie Institute for Clinical Brain Research and Department for Neurosurgery, University of Tübingen, 72074 Tübingen, Germany.; University Lille, EA7364 RADEME, CHU Lille, Institut de Genetique Medicale, F-59000 Lille, France.; Department of General Paediatrics, Division of Child Neurology and Inherited Metabolic Diseases, Centre for Paediatrics and Adolescent Medicine, University Hospital Heidelberg, 69120 Heidelberg, Germany.; Department of Pathology and Laboratory Medicine, Children’s Mercy Hospital and Clinics, Kansas City, MO 64108, USA.; Pediatric Neurology and Neurophysiology, Instituto de Neurologia de Goiania, Goiania 74210, Brazil.; Department of Medical Genetics, AP-HM, Hôpital d’Enfants de La Timone, 13005 Marseille, France.; Marseille Medical Genetics Center, Aix Marseille Univ, Inserm, U1251, Marseille, France.; Center for Fetal Diagnostics, Aarhus University Hospital, 8200 Aarhus, Denmark.; Diagnostic Imaging, Hospital for Sick Children, ON M5G 1X8, Canada.; Department of Immunology and Microbial Science, The Scripps Research Institute, CA 92037, USA.; Manchester Centre for Genomic Medicine, St Mary’s Hospital, Manchester University NHS Foundation Trust, Health Innovation Manchester, Manchester M13 9WL, UK.; Department of Neurology, Boston Children’s Hospital, Harvard Medical School, MA 02115, USA.

## Abstract

We delineate a *KMT2E* gene-related neurodevelopmental disorder based on 38 individuals in 36 families. This includes 31 distinct heterozygous variants in the *KMT2E* gene (28 ascertained from Matchmaker Exchange and 3 previously reported), and 4 individuals with chromosome 7q22.2-22.23 microdeletions encompassing the *KMT2E* gene (1 previously reported). Almost all variants occurred *de novo*, and most were truncating. Most affected individuals with protein-truncating variants presented with mild intellectual disability. One-quarter of individuals met criteria for autism. Additional common features include macrocephaly, hypotonia, functional gastrointestinal abnormalities, and a subtle facial gestalt. Epilepsy was present in about one-fifth of individuals with truncating variants, and was responsive to treatment with anti-epileptic medications in almost all. Over 70% of the individuals were male and expressivity was variable by sex, with epilepsy more common in females and autism more common in males. The four individuals with microdeletions encompassing *KMT2E* generally presented similarly to those with truncating variants, but the degree of developmental delay was greater. The group of four individuals with missense variants in *KMT2E* presented with the most severe developmental delays. Epilepsy was present in all individuals with missense variants, often manifesting as treatment-resistant infantile epileptic encephalopathy. Microcephaly was also common in this group. Haploinsufficiency versus gain-of-function or dominant negative effects specific to these missense variants in *KMT2E* may explain this divergence in phenotype, but requires independent validation. Disruptive variants in KMT2E are an under-recognized cause of neurodevelopmental abnormalities.

## Main Text

*KMT2E* encodes a member of the lysine N-methyltransferase 2 (KMT2) family. This family of enzymes plays a vital role in regulating post-translational histone methylation of histone 3 on lysine 4 (H3K4)^1^. Proper H3K4 methylation is required to maintain open chromatin states for regulation of transcription. There are at least eight known monogenic disorders impairing regulation of H3K4 methylation that present with neurodevelopmental syndromes (Table 1). In addition to these Mendelian disorders, dysregulated H3K4 methylation is believed to play a role in the pathogenesis of schizophrenia and autism^2^. *De novo* truncating variants in the *KMT2E* gene have previously been reported in three unrelated males in a large sequencing study of non-syndromic autism, but phenotypic data was limited^3–5^. In this report, we present twenty-nine additional individuals with heterozygous variants in *KMT2E* in an effort to define a KMT2E-related neurodevelopmental disorder. We also describe four individuals with chromosome 7q22.2-22.23 microdeletions encompassing *KMT2E*.

**Table 1:** Monogenic neurodevelopmental syndromes due to genes involved in regulation of H3K4 methylation

| Gene | Disorder | Inheritance | Clinical features |
| --- | --- | --- | --- |
| *KMT2A* | Wiedemann Steiner syndrome (MIM 605130) | AD | Intellectual disability, short stature, hairy elbows, dysmorphic features, autism |
| *KMT2B* | Childhood onset dystonia 28 (MIM 617284) | AD | Dystonia, developmental delay, short stature |
| *KMT2C* | Kleefstra syndrome 2 (MIM 617768) | AD | Intellectual disability, autism, epilepsy, short stature, dysmorphic features, hypotonia |
| *KMT2D* | Kabuki syndrome 1 (MIM 147920) | AD | Intellectual disability, short stature, dysmorphic features, multiple congenital anomalies, hirsutism |
| *ARX* | X-linked intellectual disability 29 (MIM 300419), Partington syndrome (MIM 309510), Proud syndrome (MIM 300004), X-linked Lissencephaly 2 (MIM 300215), Early infantile epileptic encephalopathy 1 (MIM 308350) | X-linked | Intellectual disability, dysmorphic features, brain malformations, epilepsy |
| *CUL4B* | X-linked syndromic intellectual disability 15 (Cabezas type) (MIM 300354) | X-linked | Intellectual disability, short stature, macrocephaly, epilepsy, brain malformation, behavioral abnormalities |
| *KDM5C* | X-linked syndromic intellectual disability (Claes-Jensen type) (MIM 300534) | X-linked | Intellectual disability, dysmorphic features, spasticity, short stature |

New cases were ascertained from GeneMatcher through the Matchmaker Exchange Network between September 2016 and August 2018^6,7^. All individuals were found to have variants in *KMT2E* on exome or genome sequencing. Informed consent was obtained for photographic images used in composite figure generation.

### KMT2E is constrained for protein-truncating variation in the general population

The Genome Aggregation Database (gnomAD) is a large-scale reference database with high quality, jointly processed exome or genome data from over 140,000 individuals^8^. Constraint analysis performed on the gnomAD dataset shows that *KMT2E* is a candidate haploinsufficient gene. *KMT2E* is very depleted for protein-truncating variants presumably due to negative selection, with an observed/expected ratio of 0.01 and probability of loss of function intolerance (pLI) score of 1.0.

We reviewed the 28 loss of function variants present in gnomAD v2.1. The majority of these variants are not expected to result in protein truncation for a variety of reasons including annotation artifacts (n=8), sequence errors at a simple repeat (n=5), somatic mosaicism (n=1), and a splice site rescue (n=1). Four variants are part of a complex variant in one individual that when resolved, is not expected to result in truncation. Four variants found in eight individuals in gnomAD are in the last exon; two are expected to result in truncation of the last exon and two will result in protein extension. Of note, the two protein-extension variants are located close to the variant in individual #28 (p.Val1818Alafs*48). The inheritance of this variant is unknown as the father is not available for testing, though it is not present in his mother, so this remains a variant of uncertain significance.

After review, there were five variants in gnomAD that appear to result in protein truncation. These are found in 3 males and 2 females between the ages of 30 and 70. All 5 are absent from the control only subset of gnomAD (though it should be noted that gnomAD does not contain cohorts recruited for severe, pediatric onset disease, rather contains cohorts recruited for adult onset common diseases such as cardiovascular disease and type II diabetes). By reviewing the data subsets, two appear to be from neurologic cohorts and three are from non-neuro and non-cancer cohorts. Overall, there are very few variants that are likely to result in protein truncation of *KMT2E* present in a large general population reference database.

### We ascertained thirty-eight individuals with KMT2E gene variants in association with a neurodevelopmental phenotype

Including the three previously reported cases^3–5^, thirty-four individuals from thirty-two families were ascertained with single nucleotide or indel variants in *KMT2E* and four additional individuals had copy number variants encompassing *KMT2E* (Figure 1, Table 2). The *KMT2E* variants were found to have arisen *de novo* in 26 individuals in our cohort. The variant was maternally-inherited in one of the previously reported individuals (maternal phenotype unknown)^5^. Inheritance of the variant was unknown in four families where both parents were not available for testing. In only one family was the variant found in multiple affected individuals with three affected male children; the variant was not found in their mother, and the father was not available for testing, but he was reported to have intellectual disability. Thirty variants were protein-truncating variants: twenty-four were indels, four were nonsense variants, and two were variants at essential splice sites (Figure 1A). Twenty-three of these are predicted to produce transcripts that would be subject to nonsense mediated decay. Five of the protein-truncating variants fall in the terminal exon of the gene, potentially escaping nonsense-mediated decay; three of these five variants extend the open reading frame. Only one variant was seen in two independent families (c.1776_1780delAAAGA, p.Lys593Argfs*17) in a male (individual #9) and a female (individual #10). Four of the individuals had *de novo* missense variants, three of which occur at highly conserved positions/regions of the gene (P1376S is not well conserved and serine is present in some mammalian species) (Figure 1B). There was no particular patterning or clustering of the missense variants. CADD scores are summarized in Table 2. None of the *KMT2E* variants are reported in public databases (gnomAD, Exome Variant Server, or 1000 Genomes)^8–10^.

**Figure 1:**
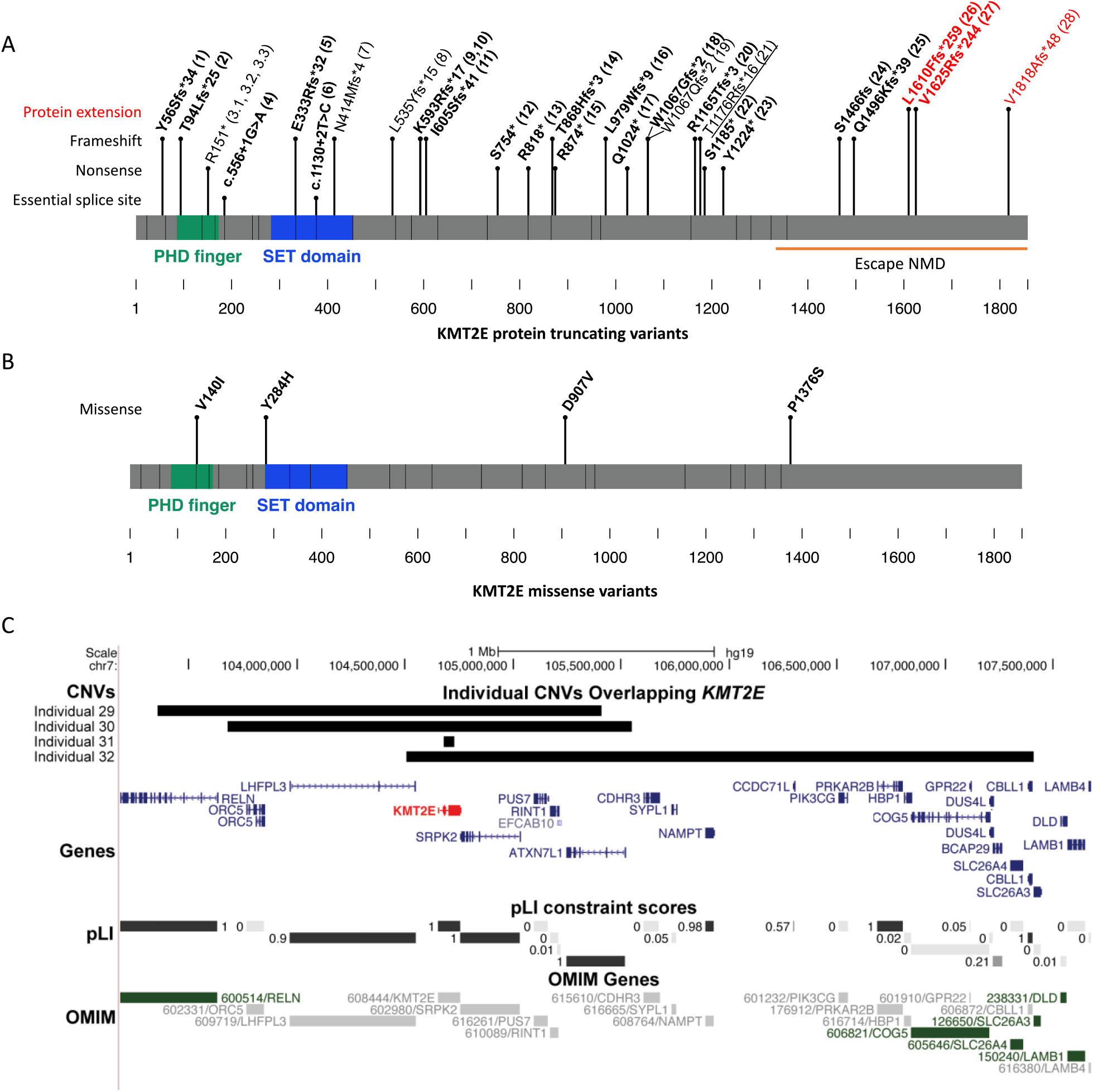
*KMT2E* variants in 38 individuals. (A) 28 protein-truncating variants in *KMT2E* identified in 30 individuals. Variants in bold are *de novo* in the proband while the underlined variant was inherited. In some cases, both parents are not available and the *de novo* status is unknown (non-bold). Variants in the last exon are predicted to escape non-sense mediated decay (individuals #24-28) while the last 3 variants (red) also result in protein extension (individuals #26-28). (B) Missense variants in *KMT2E* in individuals #33-36. (C) *De novo* deletions overlapping *KMT2E* were identified in individuals #29-32. All OMIM gene-disease associations (green) are for recessive disease.

**Table 2:**
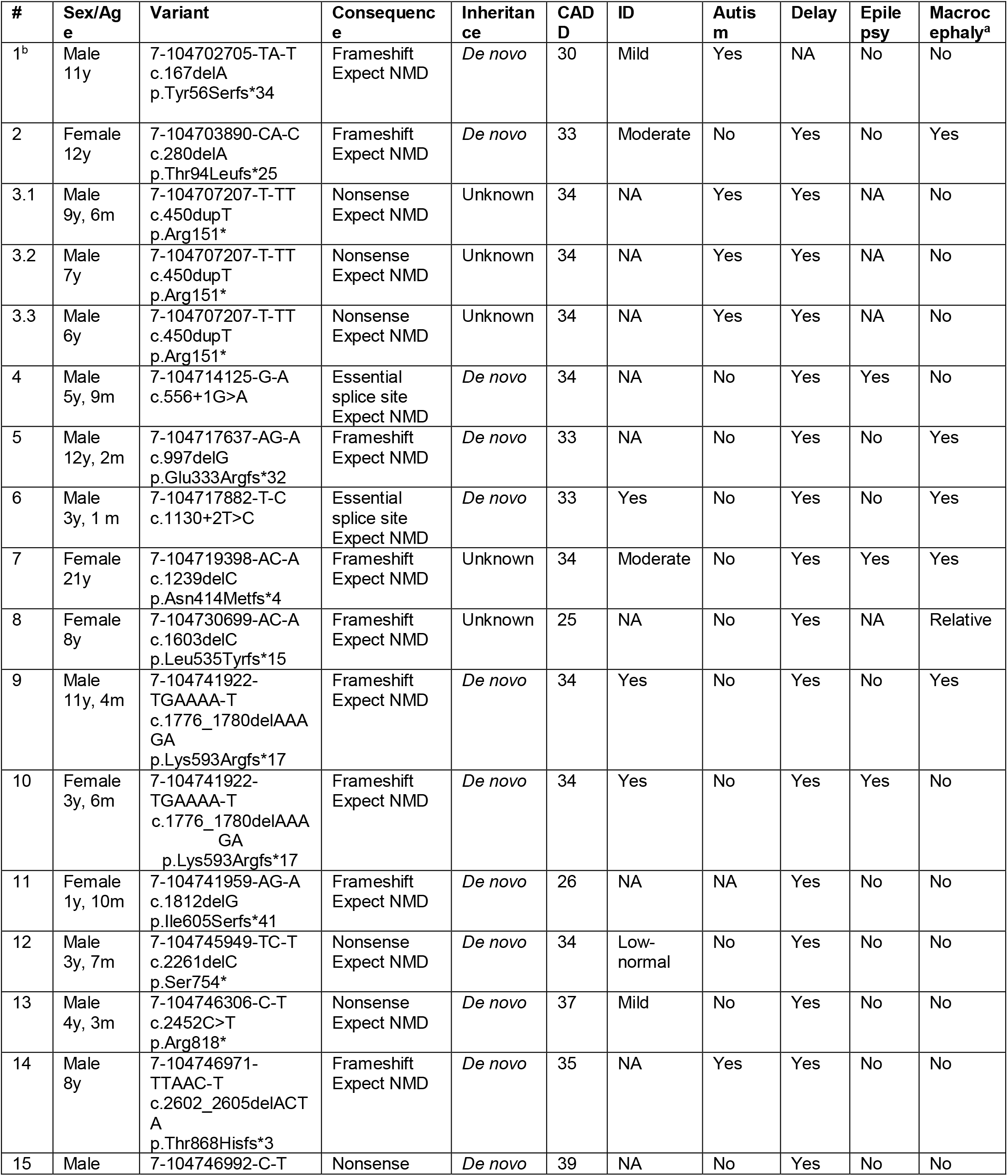
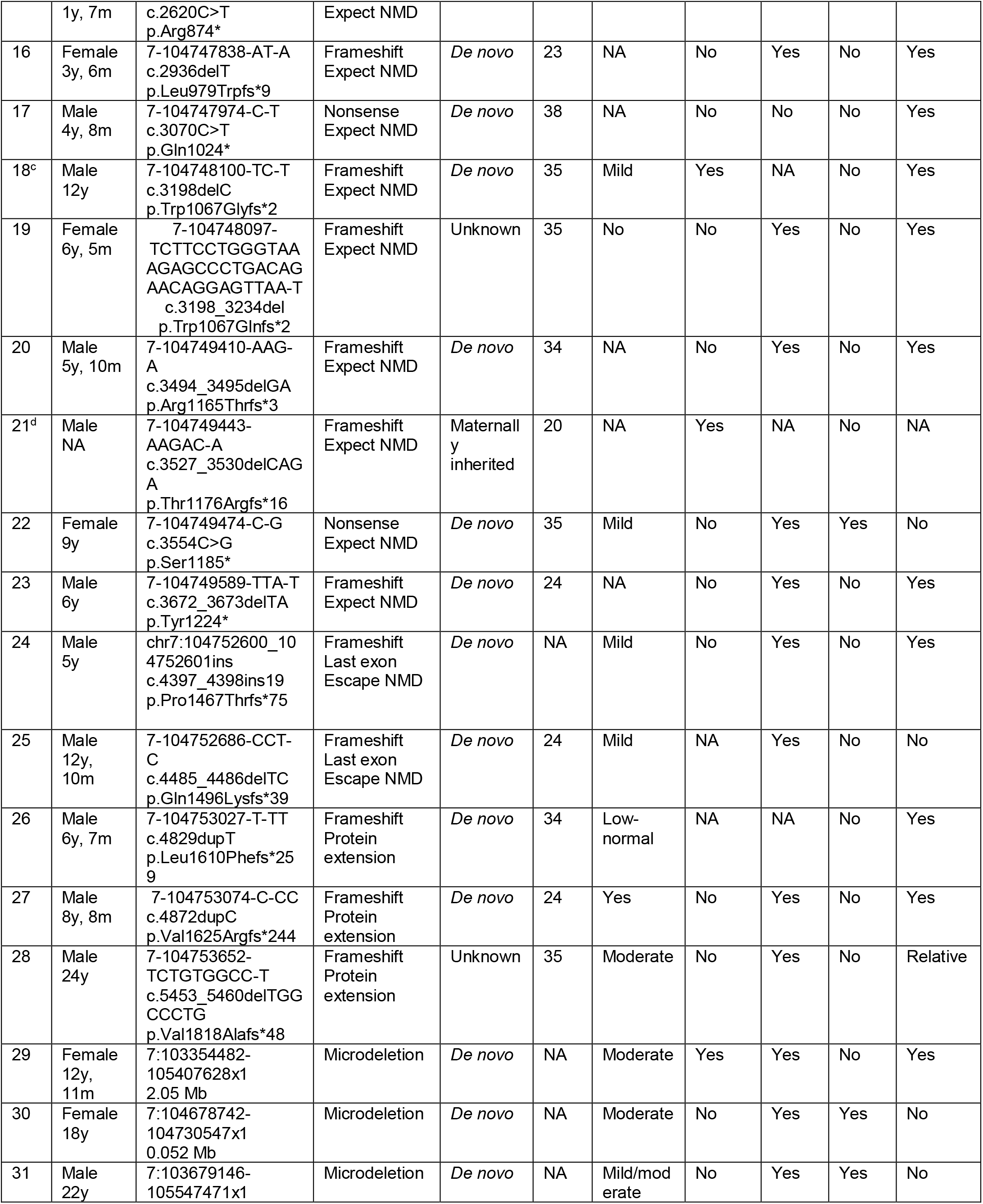
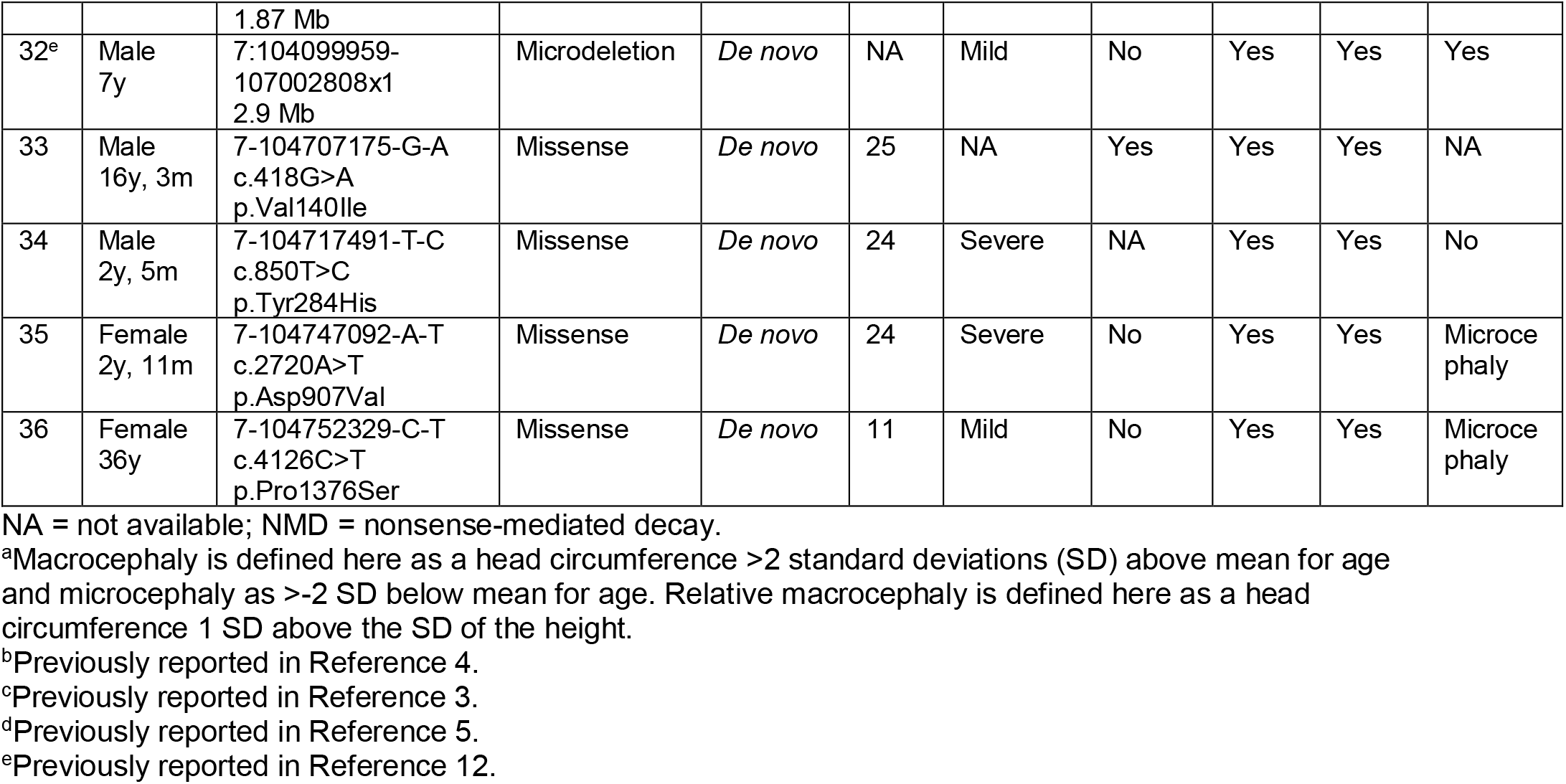
Summary of *KMT2E* Variants Found in 38 Individuals with Neuro-developmental Phenotypes

For the four individuals with chromosome microdeletions encompassing *KMT2E*, all deletions occurred *de novo*. Deletion sizes range from 0.052 to 3.2 Mb. The 0.052 Mb deletion in individual #30 involves only *KMT2E*, whereas the other three deletions include additional genes^11^. Figure 1C illustrates the genes included in these deletions. Median maternal and paternal age was 30 and 36 years, respectively. There were phenotypic differences between individuals with protein-truncating variants, missense and copy number variants, as summarized below.

### Thirty-four individuals with KMT2E Protein-truncating variants

For the individuals with protein-truncating variants in *KMT2E*, 22 were male and eight were female. Age at most recent evaluation ranged from 19 months to 24 years. Prenatal and neonatal courses were largely uncomplicated for most individuals with protein-truncating variants. One individual was born prematurely at 35 weeks. Several individuals had neonatal jaundice, one had hypoglycemia, one had sinus tachycardia, and two had neonatal feeding difficulties. Individual #10 developed respiratory arrest at fourteen hours of life and had a hypoxic-ischemic injury with typical sequelae seen on neuroimaging. She has spastic quadriplegia and epilepsy, and is not included in the subsequent paragraphs below since her acquired injury significantly influences her phenotype and is likely not representative of the disorder itself (although it cannot be excluded that the genetic disorder predisposed to the injury).

Of the remaining 29 individuals in this group (i.e. excluding individual #10), 24 had early developmental delay. The three individuals without documented developmental delay are the cases previously reported from autism studies where only limited clinical information is available^3–5^. The mean age of independent walking in this group was 20 months (range 12-48 months, Figure 2). All individuals are currently able to walk independently. Twelve of the 29 individuals have hypotonia. Individual #15 had normal initial motor development, but developed progressive spastic diplegia at 14 months of age. Neuroimaging in this individual demonstrated cerebral white matter abnormalities.

**Figure 2:**
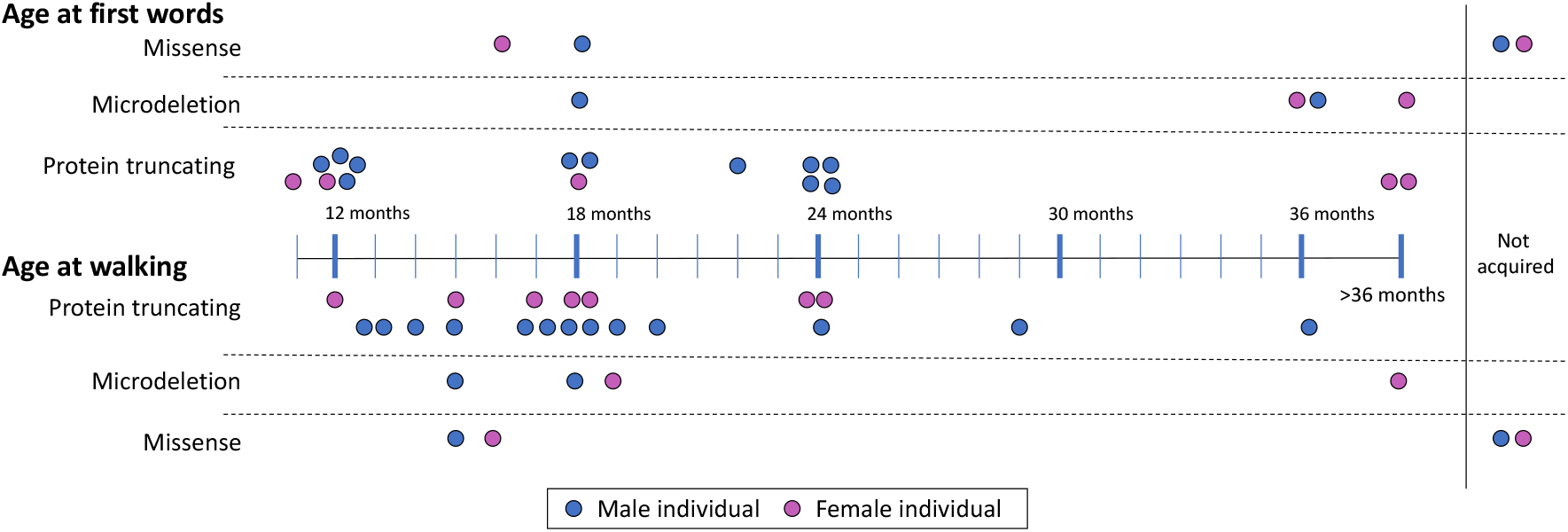
Developmental milestones in individuals with variants in *KMT2E*. Most children with protein-truncating variants acquire first words and walking by 24 months of age, though a minority are more significantly delayed. Only individual #12 who experienced a cardiac arrest and injury did not acquire these skills. A majority of individuals with microdeletion had significant delay in speech development but walked at a similar time to individuals with protein-truncating variants. For those with missense variants, those with severe infantile epilepsy had significant delays.

The mean age of acquired first word in this group was 20 months (range 12-48 months, Figure 2). Though this information is not available for all individuals, 14 (out of 17) individuals are verbal, though seven are noted to speak poorly or have articulation problems. Three of the individuals were reported to have speech regression. Intelligence quotient (IQ) data were available for only seven out of the 29 individuals: the mean IQ was 74 (range 62-98). Seven of the individuals have been diagnosed with autism. One additional individual was diagnosed with a sensory integration disorder, and another with difficulty in social interaction not meeting criteria for autism. At least two of the individuals have been diagnosed with attention-deficit/hyperactivity disorder (ADHD). Additional behavioral concerns were reported in eleven of the individuals, including stereotypies, skin picking behavior, self-injurious behavior, aggression, and anxiety.

Fourteen of the 30 individuals had macrocephaly, defined by a head circumference equal to two or more standard deviations above the mean, or 95th percentile or greater. An additional two individuals have relative macrocephaly, defined here as head circumference one standard deviation higher than the standard deviation for the height. Individual #6 also had a *de novo* pathogenic *PTEN* variant, which can also account for his macrocephaly. Other growth parameters were variable for individuals in this group, but most were in the normal range for height and weight.

Excluding individual #10 with hypoxic-ischemic injury, only four of the individuals in this group had epilepsy (two or more unprovoked seizures) (#4, #7, #8, #22); an additional patient had a history of just one seizure at eight years of age (#9). There was no consistent seizure semiology or epilepsy syndrome described across the individuals. Only one of the four individuals with epilepsy had treatment-resistant epilepsy (#7). Nineteen of the individuals had at least one brain MRI. MRI findings were normal or non-specific, with no consistent abnormalities. Thinning or partial agenesis of the corpus callosum (individuals #5, #12, #15), various cysts including pineal, epidermoid, arachnoid, ependymal (in individuals #6, #7, #8, #19, respectively), increased white matter signal (individual #8, #17), hyperintense signal in the basal ganglia (individual #10), decreased volume (individuals #5, #10, #12, #15), delayed myelination (individual #19), small areas of heterotopia (individual #20) and Chiari I malformation (individual #14).

Many of the individuals were reported to have gastrointestinal symptoms, including reflux, vomiting, or bowel motility issues; these are issues commonly seen in individuals with hypotonia. All individuals tested had normal hearing. There were no significant ophthalmological findings. There were no other recurrent health complications noted in this group. Comparing individuals with truncating variants in the terminal exon of *KMT2E* to those with earlier truncating variants, there were no clear phenotypic differences, though the number of individuals available for comparison is small.

It is notable that 22 out of the 30 individuals with protein-truncating variants were male. Additionally, the expressivity of certain aspects of the phenotype is variable between males and females (Table 3). While the rate of intellectual disability and macrocephaly were consistent, interestingly, epilepsy was seen in 43% of females and in only 5% of males (p=0.04, Fisher’s Exact test), while autism was seen in 35% of males and in none of the females (p=0.14, Fisher’s Exact test) with protein-truncating variants in *KMT2E*. It is possible that there is decreased penetrance or variable expressivity of the condition in females, leading to fewer female individuals with *de novo* protein-truncating variants coming to diagnostic attention.

**Table 3:** Summarized Phenotypes by Variant Type

| Variant type | Subset | # | Intellectual Disability | Autism | Epilepsy | Macrocephaly | Microcephaly |
| --- | --- | --- | --- | --- | --- | --- | --- |
| **Protein-truncating variants (PTVs)** | Total | 30 | 81%<br>(13/16) | 26%<br>(7/27) | 15%<br>(4/26) | 55%<br>(16/29) | 0%<br>(0/29) |
|  | Male | 22 | 82%<br>(9/11) | 35%<br>(7/20) | 5%<br>(1/19) | 52%<br>(11/21) | 0%<br>(0/21) |
|  | Female | 8 | 80%<br>(4/5) | 0%<br>(0/7) | 43%<br>(3/7) | 63%<br>(5/8) | 0%<br>(0/8) |
| **Microdeletion** | Total | 4 | 100%<br>(4/4) | 25%<br>(1/4) | 75%<br>(3/4) | 50%<br>(2/4) | 0%<br>(0/4) |
| **Missense** | Total | 4 | 100%<br>(3/3) | 33%<br>(1/3) | 100%<br>(4/4) | 0%<br>(0/3) | 66%<br>(2/3) |

### Four individuals with 7q22.2-22.3 chromosome deletions involving KMT2E

For the four individuals with *de novo* copy number variants including *KMT2E*, two were male and two were female. Age at most recent evaluation ranged from 7 to 22 years. Clinically, patients with deletions presented similarly to those with truncating variants. While the sample size is small, there appear to be more severe developmental delays in this group. Average age of first words was 34.5 months (range 18 to 48 months, Figure 2). Only two of the four individuals are verbal. Walking was delayed in all, with a range of 15-42 months. Three of the four individuals in this group have epilepsy (#30, #31, #32). Two of the four individuals in this group have macrocephaly (#29, #32).

Individual #32 has been previously reported^12^. He presented with global developmental delay, overgrowth, macrocephaly, delayed bone age, and treatment refractory generalized epilepsy. MRI of the brain demonstrated reduction of cerebral white matter, corpus callosum hypoplasia, right cerebellar hypoplasia, and an enlarged cisterna magna. Brain imaging was also performed in individuals #30 and #31. The MRI of individual #31 demonstrated global cerebral atrophy, and the MRI of individual #30 demonstrated a possible focal cortical dysplasia.

### Four individuals with de novo KMT2E missense variants

For the four individuals with *de novo* missense variants in *KMT2E*, two were male and two were female. Age at most recent evaluation ranged from 29 months to 36 years. All four of the individuals with missense variants had epilepsy. Individual #33 had five generalized tonic-clonic seizures, starting at the age of 15 years. Individuals #34, #35, and #36 all presented with infantile epileptic encephalopathy. Individual #34 developed seizures at 6 months of age, and individuals #35 and #36 both developed seizures in the neonatal period. Reported seizure semiologies include generalized tonic-clonic, tonic, atonic, myoclonic seizures, and epileptic spasms. The initial EEG in individual #35 showed burst-suppression, and subsequently evolved into hypsarrhythmia. The EEG in individual #36 also showed hypsarrhythmia. The EEG in patient #34 demonstrated background disorganization, and multifocal and generalized epileptiform discharges. All three individuals have treatment-resistant epilepsy. Individual #34 was started on the ketogenic diet at 14 months of age, which did not improve seizure control.

In our cohort, individuals with missense variants also had more severe developmental delays compared to the individuals with truncating variants. Only two of the four individuals can walk independently, and none of the individuals are verbal at most recent follow-up (Figure 2). Two of the four individuals in this category have microcephaly, and the other two are normocephalic. Three of these individuals had a brain MRI: one individual had delayed myelination, one had cerebral atrophy, and one had an incidental abnormality in the right cerebral peduncle.

### Facial analysis

Comparison of the facial features of eleven of the individuals in our cohort suggests some commonalities, including macrocephaly, dolichocephaly, high forehead, deep-set eyes, periorbital fullness, prominent cheeks, and prominent nasolabial folds (Figure 3). Utilizing Face2Gene (FDNA, Inc., Boston, MA) facial recognition software, we created a composite image from frontal photographs of these 11 individuals (excluding individual #30 with glasses) to represent the common facial gestalt.

**Figure 3:**
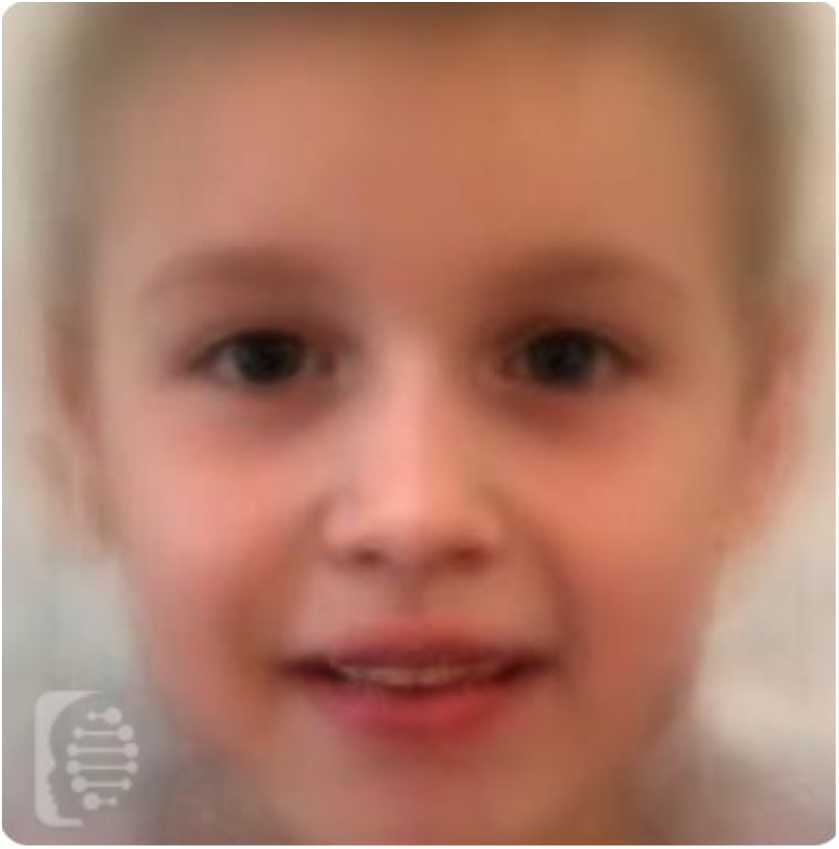
Composite photo from Face2Gene. Photos of 11 individuals were used in this analysis (eight with PTVs, two with microdeletions and 1 with a missense variant).

*KMT2E* encodes a histone methyltransferase protein, a transcriptional regulator reported to play key roles in diverse biological processes, including cell cycle progression, maintenance of genomic stability, adult hematopoiesis, and spermatogenesis. The gene is highly expressed in the brain, particularly during fetal development^4^. *KMT2E* appears to be distinct from other members of the KMT2 family. Most KMT2 proteins contain an enzymatically active SET domain that possesses methyltransferase function^2,13^. While the KMT2E protein contains a SET domain, it is different in sequence and location within the protein than other members of the KMT2 family, and studies suggest that it may lack intrinsic methyltransferase activity^14^. However, the SET domain is still highly conserved in KMT2E, and it has been proposed that KMT2E may have an indirect effect on H3K4 methylation, possibly through transcriptional regulation of additional histone modifying enzymes. Most members of the KMT2 family contain multiple PHD finger domains that function as H3K4 methylation readers. In contrast, *KMT2E* contains a single PHD finger domain. PHD fingers typically bind to specific epigenetic histone marks in order to recruit transcription factors and nucleosome-associated complexes to chromatin. Finally, while most members of the KMT2 family function as global activators of open chromatin, *KMT2E* is believed to be a repressor, although the precise mechanisms involved in *KMT2E* regulation of gene transcription have not yet been elucidated^15^.

The individuals with protein-truncating *KMT2E* variants in our cohort present with syndromic intellectual disability. Most individuals are functioning in the low-normal to mild intellectual disability range. Seven of the individuals (including of the three previously reported individuals^3–5^) have also been formally diagnosed with autism. There appears to be a subtle common facial gestalt amongst the individuals whose images were available for review. Additional features, albeit not obligate or specific, include macrocephaly, hypotonia, and GI dysmotility. Neuroimaging is normal or non-specific. Epilepsy was not common among the individuals with protein-truncating variants. There were no significant phenotypic differences between individuals with truncating variants in the terminal exon of the gene and earlier truncating variants, suggesting a probable common pathophysiology of haploinsufficiency.

In contrast to the individuals with protein-truncating variants, the individuals we report with missense variants all have epilepsy. Three of these individuals fall in the category of an infantile-onset epileptic encephalopathy. In addition, these individuals have more severe developmental delays, and two have microcephaly. We hypothesize that the phenotype of epileptic encephalopathy may be variant specific, and may relate to an alternate mechanism such as gain-of-function or dominant negative effect. Recently distinct developmental disorder phenotypes have been identified to result from PTVs and missense variants in the same gene^16,17^. Additional cases and further functional studies are required to clarify this.

Overall, the individuals with chromosome 7q22.2-22.3 microdeletions encompassing *KMT2E* presented similarly to those with truncating variants, further supporting haploinsufficiency as the disease mechanism. While the sample size was small, these individuals appeared to have more severe developmental delays compared to those individuals with truncating variants, which is likely explained by the influence of additional genes included in their deletions. The 7q22.2-22.3 region contains multiple additional genes involved in the regulation of the cell cycle, including *SRPK2, RINT1*, and *LHFPL*3^12^. In particular, the *SRPK2* gene is also high constrained in the gnomAD database (pLI of 1.0) and is expressed in the central nervous system. The *SRPK2* gene encodes a cell-cycle regulated protein kinase that phosphorylates serine/arginine domain-containing proteins and modulates pre-mRNA splicing in neurons^18^.

Several *Kmt2e (Mll5)* deficiency mouse models have been created and characterized^15,19–22^. These mice present with growth restriction and increased mortality, as well as impaired hematopoiesis. A neurological phenotype in these mice has not been reported. Both homozygous and heterozygous loss of *Kmt2e* in mice results in DNA damage and elevated levels of reactive oxygen species (ROS)^22^. The cellular effects were effectively reversed by supplementation with the glutathione precursor, N-acetylcysteine (NAC)^22^. This has interesting therapeutic implications in humans, since NAC supplementation has been used to treat glutathione depletion in acetaminophen overdose as well as rare inborn errors of metabolism associated with increased free radical damage. Further studies are required to establish whether humans haploinsufficient for *KMT2E* are also vulnerable to increased ROS, and whether there may be a benefit in treating with NAC or other antioxidants.

In this report, we define a KMT2E-related neurodevelopmental disorder, which adds to the growing list of KMT2 gene family disorders. Most individuals with protein-truncating variants appear to present with generally mild developmental delay/intellectual disability. Autism is also relatively common. Additional common, but not obligate, features include relative macrocephaly, hypotonia, and functional gastrointestinal disturbances. There appears to be a subtle facial gestalt. Epilepsy was not common amongst individuals with protein-truncating variants. We suspect haploinsufficiency as the disease mechanism. The similar phenotype seen in individuals with microdeletions of this region is consistent with this hypothesis. In contrast, individuals with missense variants all presented with epilepsy, including infantile-onset epileptic encephalopathy, and more severe developmental delays. Variant specific alterations in *KMT2E* function, possibly even gain-of-function, may explain this divergence in phenotype. Further studies are required to further understand genotype-phenotype correlation. There is no established therapy for KMT2E-related disorders, although based on animal data, there may be a role for N-acetylcysteine or other antioxidant treatments.

## Acknowledgements

We thank the families for participating in this study, GeneMatcher, and the gnomAD team for the Genome Aggregation Database. We are thankful to the Deciphering Developmental Disorders (DDD) study for the invaluable collaboration. We are appreciative to Eric Minikel for sharing the R code that was adapted for Figure 1. Face2Gene software was used to generate Figure 3 and we appreciate guidance from Nicole Fleischer for using this software. We appreciate Steven Harrison’s guidance for variant curation and ClinVar submission.

Support was provided to AHOL by the National Institute of Health funded by the National Child Health and Development Institute (K12 HD052896) and by Boston Children’s Hospital Office of Faculty Development BTREC Faculty Career Development Fellowship; to LSP through the Broad Center for Mendelian Genomics funded by the National Human Genome Research Institute (UM1 HG008900); to VF by CONICYT, Chile’s National Commission for Scientific and Technological Research (grant number 72160007); to SS by a grant from the Dietmar-Hopp-Stiftung (1DH1813319); and to JZ by a Skaggs-Oxford Scholarship. JJD is supported by the Mogford Campbell family chair and by the Canadian Institute for Health Research (363863). JCT is funded by the Health Innovation Challenge Fund (R6-388 / WT 100127), a parallel funding partnership between the Wellcome and the Department of Health. The DDD Study (Cambridge South REC approval 10/H0305/83 and the Republic of Ireland REC GEN/284/12) presents independent research commissioned by the Health Innovation Challenge Fund (grant number HICF-1009-003), a parallel funding partnership between the Wellcome and the Department of Health, and the Wellcome Sanger Institute (grant number WT098051). The research team acknowledges the support of the National Institute for Health Research, through the Comprehensive Clinical Research Network. This study makes use of DECIPHER (http://decipher.sanger.ac.uk), which is funded by the Wellcome. The views expressed in this publication are those of the authors and not necessarily those of the Wellcome or the Department of Health. YW and IH were funded by the German Research Society (DFG WE4896/3-1, WE4896/4-1, HE5415/3-1, HE5415/5-1, HE5415/6-1, HE5415/7-1). IH was also supported by intramural funds of the Children’s Hospital of Philadelphia and the University of Kiel.

## References

1. Shen, E., Shulha, H., Weng, Z., and Akbarian, S. (2014). Regulation of histone H3K4 methylation in brain development and disease. Philos. Trans. R. Soc. Lond. B Biol. Sci. 369,.

2. Faundes, V., Newman, W.G., Bernardini, L., Canham, N., Clayton-Smith, J., Dallapiccola, B., Davies, S.J., Demos, M.K., Goldman, A., Gill, H., et al. (2018). Histone Lysine Methylases and Demethylases in the Landscape of Human Developmental Disorders. Am. J. Hum. Genet. 102, 175–187.

3. Iossifov, I., Ronemus, M., Levy, D., Wang, Z., Hakker, I., Rosenbaum, J., Yamrom, B., Lee, Y.-H., Narzisi, G., Leotta, A., et al. (2012). De novo gene disruptions in children on the autistic spectrum. Neuron 74, 285–299.

4. Dong, S., Walker, M.F., Carriero, N.J., DiCola, M., Willsey, A.J., Ye, A.Y., Waqar, Z., Gonzalez, L.E., Overton, J.D., Frahm, S., et al. (2014). De novo insertions and deletions of predominantly paternal origin are associated with autism spectrum disorder. Cell Rep. 9, 16–23.

5. Wang, T., Guo, H., Xiong, B., Stessman, H.A.F., Wu, H., Coe, B.P., Turner, T.N., Liu, Y., Zhao, W., Hoekzema, K., et al. (2016). De novo genic mutations among a Chinese autism spectrum disorder cohort. Nat. Commun. 7, 13316.

6. Sobreira, N., Schiettecatte, F., Valle, D., and Hamosh, A. (2015). GeneMatcher: a matching tool for connecting investigators with an interest in the same gene. Hum. Mutat. 36, 928–930.

7. Philippakis, A.A., Azzariti, D.R., Beltran, S., Brookes, A.J., Brownstein, C.A., Brudno, M., Brunner, H.G., Buske, O.J., Carey, K., Doll, C., et al. (2015). The Matchmaker Exchange: a platform for rare disease gene discovery. Hum. Mutat. 36, 915–921.

8. Lek, M., Karczewski, K.J., Minikel, E.V., Samocha, K.E., Banks, E., Fennell, T., O’Donnell-Luria, A.H., Ware, J.S., Hill, A.J., Cummings, B.B., et al. (2016). Analysis of protein-coding genetic variation in 60,706 humans. Nature 536, 285–291.

9. Tennessen, J.A., Bigham, A.W., O’Connor, T.D., Fu, W., Kenny, E.E., Gravel, S., McGee, S., Do, R., Liu, X., Jun, G., et al. (2012). Evolution and functional impact of rare coding variation from deep sequencing of human exomes. Science 337, 64–69.

10. 1000 Genomes Project Consortium, Auton, A., Brooks, L.D., Durbin, R.M., Garrison, E.P., Kang, H.M., Korbel, J.O., Marchini, J.L., McCarthy, S., McVean, G.A., et al. (2015). A global reference for human genetic variation. Nature 526, 68–74.

11. Kent, W.J., Sugnet, C.W., Furey, T.S., Roskin, K.M., Pringle, T.H., Zahler, A.M., and Haussler, D. (2002). The human genome browser at UCSC. Genome Res. 12, 996–1006.

12. Uliana, V., Grosso, S., Cioni, M., Ariani, F., Papa, F.T., Tamburello, S., Rossi, E., Katzaki, E., Mucciolo, M., Marozza, A., et al. (2010). 3.2 Mb microdeletion in chromosome 7 bands q22.2-q22.3 associated with overgrowth and delayed bone age. Eur. J. Med. Genet. 53, 168–170.

13. Rao, R.C., and Dou, Y. (2015). Hijacked in cancer: the KMT2 (MLL) family of methyltransferases. Nat. Rev. Cancer 15, 334–346.

14. Mas-Y-Mas, S., Barbon, M., Teyssier, C., Déméné, H., Carvalho, J.E., Bird, L.E., Lebedev, A., Fattori, J., Schubert, M., Dumas, C., et al. (2016). The Human Mixed Lineage Leukemia 5 (MLL5), a Sequentially and Structurally Divergent SET Domain-Containing Protein with No Intrinsic Catalytic Activity. PLoS One 11, e0165139.

15. Zhang, X., Novera, W., Zhang, Y., and Deng, L.-W. (2017). MLL5 (KMT2E): structure, function, and clinical relevance. Cell. Mol. Life Sci.

16. Rivière, J.-B., van Bon, B.W.M., Hoischen, A., Kholmanskikh, S.S., O’Roak, B.J., Gilissen, C., Gijsen, S., Sullivan, C.T., Christian, S.L., Abdul-Rahman, O.A., et al. (2012). De novo mutations in the actin genes ACTB and ACTG1 cause Baraitser-Winter syndrome. Nat. Genet. 44, 440–444, S1–S2.

17. Cuvertino, S., Stuart, H.M., Chandler, K.E., Roberts, N.A., Armstrong, R., Bernardini, L., Bhaskar, S., Callewaert, B., Clayton-Smith, J., Davalillo, C.H., et al. (2017). ACTB Loss-of-Function Mutations Result in a Pleiotropic Developmental Disorder. Am. J. Hum. Genet. 101, 1021–1033.

18. Hong, Y., Chan, C.B., Kwon, I.-S., Li, X., Song, M., Lee, H.-P., Liu, X., Sompol, P., Jin, P., Lee, H.-G., et al. (2012). SRPK2 phosphorylates tau and mediates the cognitive defects in Alzheimer’s disease. J. Neurosci. 32, 17262–17272.

19. Heuser, M., Yap, D.B., Leung, M., de Algara, T.R., Tafech, A., McKinney, S., Dixon, J., Thresher, R., Colledge, B., Carlton, M., et al. (2009). Loss of MLL5 results in pleiotropic hematopoietic defects, reduced neutrophil immune function, and extreme sensitivity to DNA demethylation. Blood 113, 1432–1443.

20. Madan, V., Madan, B., Brykczynska, U., Zilbermann, F., Hogeveen, K., Döhner, K., Döhner, H., Weber, O., Blum, C., Rodewald, H.-R., et al. (2009). Impaired function of primitive hematopoietic cells in mice lacking the Mixed-Lineage-Leukemia homolog MLL5. Blood 113, 1444–1454.

21. Yap, D.B., Walker, D.C., Prentice, L.M., McKinney, S., Turashvili, G., Mooslehner-Allen, K., de Algara, T.R., Fee, J., de Tassigny, X.D., Colledge, W.H., et al. (2011). Mll5 is required for normal spermatogenesis. PLoS One 6, e27127.

22. Tasdogan, A., Kumar, S., Allies, G., Bausinger, J., Beckel, F., Hofemeister, H., Mulaw, M., Madan, V., Scharfetter-Kochanek, K., Feuring-Buske, M., et al. (2016). DNA Damage-Induced HSPC Malfunction Depends on ROS Accumulation Downstream of IFN-1 Signaling and Bid Mobilization. Cell Stem Cell 19, 752–767.

